# Resolving ESCRT-III spirals at the intercellular bridge of dividing cells using 3D STORM imaging

**DOI:** 10.1101/194613

**Authors:** Inna Goliand, Tali Dadosh, Natalie Elia

## Abstract

The ESCRT machinery mediates membrane fission in a verity of processes in cells. According to the proposed mechanism, ESCRT-III proteins drive membrane fission by assembling into helical filaments on membranes. Yet, ESCRT-III filaments have never been directly visualized in a cellular process that utilizes this machinery for its function. Here we used 3D STORM imaging of endogenous ESCRT-III component IST1, to describe the structural organization of ESCRT-III during mammalian cytokinetic abscission. Using this approach, ESCRT-III ring and spiral assemblies were resolved at the intercellular tube of cells undergoing abscission. Characterization of these structures indicates the ESCRT-III helical filament undergoes remodeling during abscission. This work provides the first evidence that ESCRT-III proteins assemble into helical filaments in physiological context, indicating that the ESCRT-III machine indeed derives its contractile activity through spiral assemblies. Moreover, it provides new structural information on ESCRT-III filaments, which raise new mechanistic scenarios for ESCRT driven membrane constriction.

## INTRODUCTION

The endosomal sorting complex required for transport (ESCRT) is an evolutionary conserved machine that drives membrane constriction and fission in a variety of processes in cells (Hurley, 2015). These processes include multivesicular body biogenesis, viral budding, nuclear envelope reformation, nuclear pore complex quality control and abscission of the intercellular bridge during cytokinesis. The ESCRT machinery is composed of five different subfamilies: ESCRT 0, I, II, III and the AAA ATPase VPS4. Early ESCRT components (ESCRT 0, I, II) are essential for recruiting cytosolic ESCRT-III monomers to the membrane. Polymerization of membrane bound ESCRT-III into filaments followed by disassembly of the filaments, mediated by the ATPase activity of VPS4, is thought to drive membrane constriction and fission (Hurley, 2015).

Over 10 different ESCRT III proteins were identified in the mammalian genome (CHMP1A, B; CHMP2A, B; CHMP3; CHMP4A, B, C; CHMP5; CHMP6; CHMP7 and IST1). Exactly how these monomers assemble in cell to drive membrane fission is currently unknown. In vitro, ESCRT III proteins were shown to assemble into tubes, spirals, cones and coils composing different diameters (Chiaruttini et al., 2015; Lata et al., 2008; McCullough et al., 2015). In cells, over expression of the ESCRT III protein CHMP4B induced the formation of cortical filaments and spirals on the inner side of the plasma membrane, and over expression of CHMP2B induced plasma membrane tubulations (Bodon et al., 2011; Cashikar et al., 2014; Hanson et al., 2008). ESCRT III proteins localized to these, artificially ESCRT induced, plasma membrane structures. Although these results indicate that ESCRT III proteins can form helical filaments on cellular membranes, such filaments have not been directly observed in a physiologically relevant context; i.e. in a cellular process that utilizes the ESCRT complex to drive membrane fission. Therefore, whether ESCRT-III forms helical filaments to drive membrane fission in cells could not be determined and consequently, the underlying biophysical mechanism for ESCRT mediated membrane fission remained unknown.

In cytokinetic abscission, a ∼1 μm diameter membrane tube (called the intercellular bridge) is constricted and cut in an ESCRT dependent fashion, giving rise to two independent daughter cells (supplementary Figure 1). Using Structured Illumination Microscopy (SIM), ESCRT based cortical rings were seen at the rims of the dark zone, a dense structure located at the center of the bridge. In late stages of abscission, ESCRT-III proteins were additionally found in the abscission sites, located peripherally along the bridge. Using cryo-electron-tomography (cryo-TEM) and cryo-soft-X-ray-tomography (cryo-SXT), cortical helical filaments were resolved in the region between the central dark zone and the peripheral abscission site (Guizetti et al., 2011; Sherman et al., 2016). Whether these filaments are ESCRT based has not determined. Based on these findings several models for ESCRT-mediated abscission were proposed (Elia et al., 2012; Elia et al., 2013; Guizetti and Gerlich, 2012). However, direct visualization of ESCRT spirals during abscission was needed in order to support the models.

## RESULTS AND DISCUSSION

In an attempt to directly visualize ESCRT-III assemblies at different stages of abscission we used 3D Stochastic Optical Reconstruction Microscopy (STORM). To avoid over expression artifacts we immunolabeled the ESCRT-III component IST1 in Hela cells that were synchronized to enrich the percentage of cells undergoing abscission. Microtubule staining was performed for determining abscission progression as previously described (Elia et al., 2011; Gershony et al., 2017). In bridges at early stages, IST1 was found in two large diameter cortical rings residing at the rims of the dark zone (Figure 1A, ring diameter 1.09±0.26 μm); consistent with previous data obtained for other ESCRT III proteins (Elia et al., 2011; Goliand et al., 2014).

**Figure 1:**
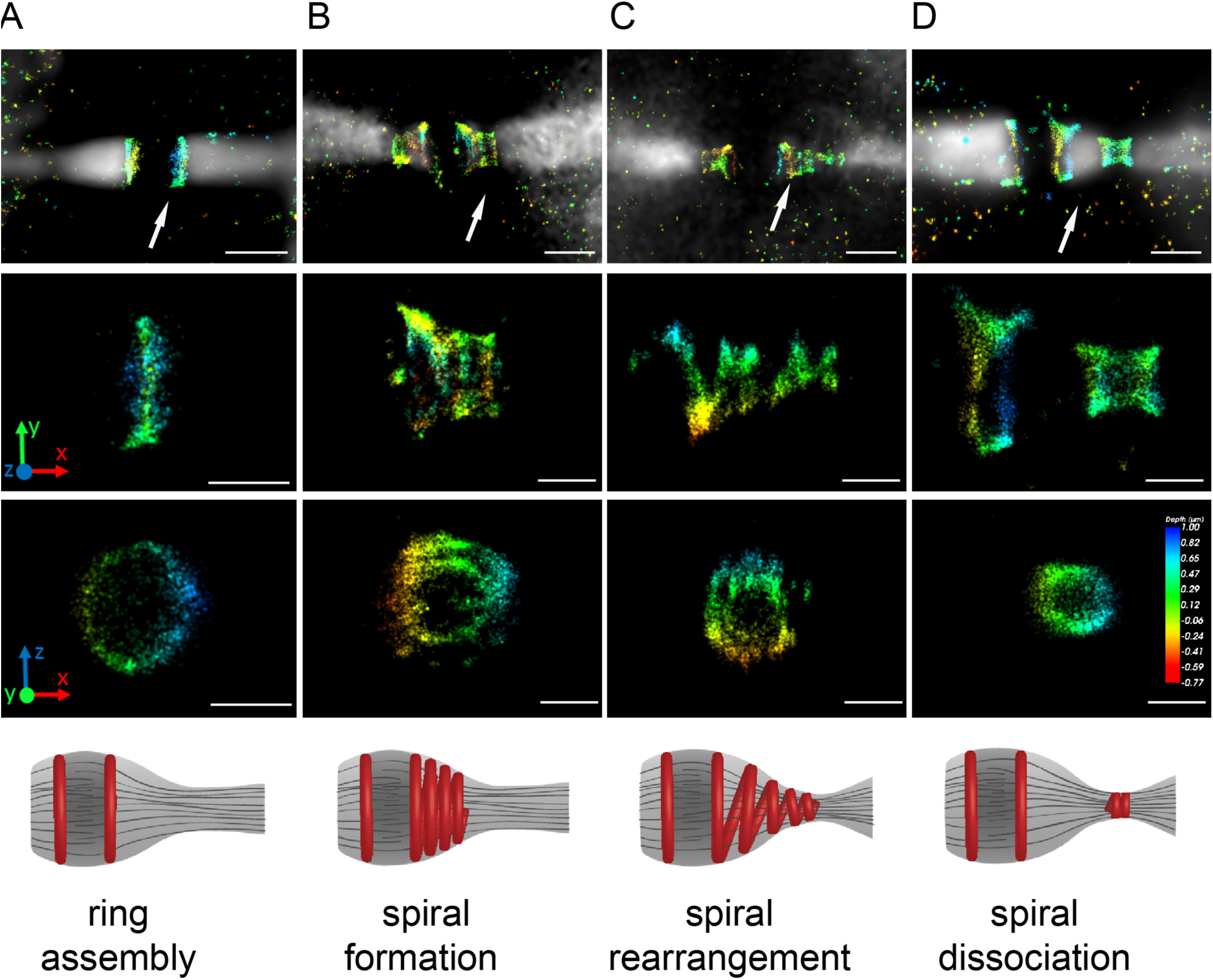
Spatial organization of the ESCRT-III protein, IST1 in the intercellular bridge of dividing cells. HeLa cells were synchronized to enrich the population of cells in late stages of cytokinesis, fixed, stained with anti–α-tubulin and anti-IST1 and imaged using the Vutara 3D STORM system. (A-D) representative images of cells at different stages of abscission (A-D, early to late, respectively), as determined based on microtubule morphology at the intercellular bridge. Microtubules were imaged in wide-field, IST1 was imaged in 3D STORM. Top Panel: a 3D-STORM image of IST1 (color coded by depth) overlaed on top of the microtubule image (white) showing the overall organization of IST1 along the intercellular bridge. The unstained area in the microtubule staining is the dense dark zone compartment of the intercellular bridge. bar, 1 μm. Second and third panels: enlarged images of IST1 structures (depicted in arrows, top panel). Images in third panel are 90° rotated. For representation reasons, the last third of the IST1 structure shown in C was omitted from the rotated image. (A) A large diameter ring is observed on each sides of the bridge (ring diameter 1.09±0.26 μm) n=31 obtained from 29 cells. (B) Accumulation of large diameter rings at the bridge (see also supplementary video 1). Largest ring diameter, 0.95±0.19 μm, smallest ring diameter 0.67±0.19 μm. n=11 obtained from 11 cells. (C) IST1 spiral (see also supplementary video 2). Largest ring diameter, 0.73±0.21 μm, smallest ring diameter 0.35±0.09 μm. n=6 obtained from 6 cells (D) A large diameter ring located at the rims of the dark zone (diameter 1±0.2 μm) and two smaller rings located at the abscission sites (diameter 0.42±0.08 μm). n=9 obtained from 6 cells. bar, 0.5 μm. Bottom panel: illustrations of the structural information presented in the panels above (microtubules, grey; ESCRT-III, red). Text denotes a proposed explanation for ESCRT-III assembly and remodeling during cytokinetic abscission based on structural information provided in the image. In short, ESCRT-III assembles at the rims of the dark zone to form a cortical ring that is stabilized by interactions with early ESCRT components (A, ring assembly). This step is followed by polymerization of ESCRT-III into a large diameter cortical spiral (B, spiral formation). Acute deformation of the spiral then leads to membrane constriction (C, spiral rearrangement). Finally, the spiral relaxes and breaks into two structures. A large diameter cortical ring held at the rims of the dark zone and a small two rings structure located at the abscission site (D, spiral dissociation).

At late stages, IST1 distributed peripherally along the bridge, forming an array of large diameter rings (Figure 1B). The rings complex was about 800 nm long and was composed of parallel rings. We were able to resolve between 2 to 4 large diameter rings in each complex. We measured an average diameter of 0.95±0.19 μm for the ring adjacent dark zone and an average diameter 0.67±0.19 μm, for the most peripheral ring. The distance between one ring to the other was 0.28±0.08 μm. In some regions, we were able to resolve connections between the rings, suggesting that they are part of a continuous spiral (supplementary video 1). It is worth noting that the inability to resolve the connectivity between some of the rings may result from the packed geometry of the spiral at this stage. In later stages, a clear helical IST1 filament was observed along the intercellular bridge. This filament stretched all the way from the initial ring assembly site, at the center of the bridge, to the site of abscission, located almost 1 micron away. In contrast to the parallel ring array observed in earlier stages, the ESCRT-III filament adopted a non-uniform helical shape composed of different diameters and pitch angles (Figure 1C, supplementary video 2). The helix adopted smaller diameters as it approached the abscission site. We measured the largest diameter for this helix at its site of origin (0.73±0.21 μm) and the smallest diameter at the abscission site (0.35±0.09 μm) (Figure 1C). Connections between the turns of the helix were clearly apparent, indicating continuity in the structure and strongly supporting a spiral structure. In final stages of abscission, IST1 localized in two pools, similarly to what we have observed for other ESCRT III proteins using SIM (Elia et al., 2011; Goliand et al., 2014). A large diameter cortical ring located at the rims of the dark zone (Figure 1D, diameter 1±0.2 μm), and a peripheral pool located at the abscission site. We consistently resolved two small diameter rings at the peripheral pool (Figure 1D, diameter 0.42±0.08 μm), indicating a high ordered organization for ESCRT-III at the abscission site.

The structural information described above is highly consistent with previous observations obtained for ESCRT-III in cytokinesis using SIM, indicating that endogenous IST1 staining represents the structural organization of ESCRT-III in abscission. Importantly, by increasing the spatial resolution in 3D and by systematically imaging cells at different stages of abscission, we were able to resolve new intermediates in ESCRT-III organization at the intercellular bridge. We show, for the first time, that ESCRT-III proteins assemble into helical filaments in a physiological process that utilizes the ESCRT complex for membrane fission. This data strongly suggest that the helical filaments previously observed at the intercellular bridge using cryo-TEM and cryo-SXT, are ESCRT based. Taken together, our data indicate that in accordance with previous in vitro data and data obtained in cells, the ESCRT-III machine mediates membrane constriction by forming transient helical filaments on the inner side of the plasma membrane.

In light of this new structural data and based on the computational model we have previously suggested for ESCRT mediated abscission (Elia et al., 2012), we propose the following scenario for ESCRT mediated abscission. ESCRT-III assembly begins with the formation of a large diameter ring, which then polymerizes into a large diameter filament. This large diameter filament accumulates stress as it grows peripherally. Lateral interactions between the turns of the packed helix or internal pulling forces from inside the tube may stabilize the unstable large diameter spiral at this stage. This step is followed by relaxation of the spiral to its preferred diameter of about 50 nm. However, because the spiral is on one side held by the large diameter ring, which is anchored to the packed dark zone region via interactions with early ESCRT components, it adopts a distorted conformation that is composed of varying diameters and pitch angles. Finally, the deformed spiral breaks into two parts: one part stays attached to the large diameter ring at the center of the bridge and the second part slides to the abscission side. Because the spiral is attached to the membrane these events leads to acute constriction of the tube, achieving a minimal tube diameter at the abscission site. Final abscission then occurs presumably according to our previously published model (Elia et al., 2012).

## MATERIALS AND METHODS

### Cell culture

HeLa cells were grown in DMEM supplemented with 10% fetal bovine serum (FBS), 2 mM glutamine, 10,000 U/ml penicillin and 10 mg/ml streptomycin in a cell culture incubator (37°C and 5% CO_2_).

### Cell culture synchronization by double thymidine block

To enrich the population of dividing cells, HeLa cells were subjected to double thymidine block as described in (Bostock et al., 1971). In short, cells were plated at 10% confluency on coverslips attached to petri dish (Ibidi, Martinsried, Germany) and incubated with growth medium containing 2 mM thymidine (SIGMA, T1895-1G). 18 hours later, thymidine was washed three times with PBS and cells were incubated for 9 hours in growth medium. Then, cells were subjected to a second thymidine block (2 mM) for 15 hours. To release cells from G1/S arrest, thymidine was washed three times with PBS and cells were incubated in fresh warm growth medium. Cells were fixed 10.5 hours after release from thymidine block to maximize the population of cells undergoing abscission.

### Immunostaining

Cells were washed with PBS solution (warmed to 37°C) and fixed with fixation buffer 3% PFA + 0.1% glutaraldehyde diluted in PBS (warmed to 37°C) for 15 minutes at room temperature. Then cells were washed with PBS and permeabilized and blocked with 0.2% Triton X-100 with 3% BSA for 30 minutes. All cells were stained with monoclonal anti α-tubulin antibodies (dilution 1:1000) (Sigma, T6199) and polyclonal anti IST1 antibody (dilution 1:50) (Proteintech, 51002-1-AP). Afterwards cells were washed with wash buffer 0.05% Triton X-100 in PBS. Cells were then subjected to a secondary antibody staining using Alexa Fluor 488 or Alexa Fluor 647 anti-mouse or anti-rabbit secondary antibodies, respectively (Life Technologies) (dilution 1:1500). All antibodies were diluted in 1% BSA + 0.2% Triton X-100 in PBS. After incubation with the secondary antibody cells were washed again with wash buffer containing 0.05% Triton X-100 in PBS. Last, cells were post fixed with fixation buffer 3% PFA +0.1% glutaraldehyde (10 minutes, room temperature) and subsequently washed with PBS.

### STORM imaging

Images were collected on Vutara SR200 STORM microscope based on the single-molecule localization biplane technology, using 60x water immersion objective (1.2 NA). Alexa Fluor 647 was excited using 647 nm laser (5 kW/cm^2^) acquiring 12,000 frames per image with acquisition time of 20 msec per frame. Imaging was performed in the presence of imaging buffer (7 μM glucose oxidase (Sigma), 56 nM catalase (Sigma), 5 mM cysteamine (Sigma), 50 mM Tris, 10 mM NaCl, 10% glucose, pH 8).

### Data Analysis

Collected particles were subjected to thresholding. Threshold value is defined in Vutara as standard deviations above the frame background value, which is determined based on the mean value of the border pixels in each frame. We typically set threshold value to 5 in our collected datasets. Particles were localized in three dimensions based on bead calibration. If needed, drift was corrected manually. Images were exported with the following visualization parameters: point splatting view, 20 nm particle size, opacity 0.03 and colorcoded by depth.

The data was subjected to de-noise and accuracy filters in the Vutara software. The denoising filter eliminates scattered points based on a nearest neighbor algorithm. Localization accuracy is calculated in Vutara following the guideline reported by Thompson et al. (Thompson et al., 2002). We obtained lateral localization accuracy of 7.2 ±0.6 in our measurements. To estimate localization accuracy in the axial dimension, we measured 1000 frames of sub diffraction limit 100nm beads (ThermoFisher, T7279) using 647 nm laser excitation. The excitation was chosen to fit the typical signal that we observed in our experiments (max counts of 5000). Fitting the histogram results, the estimated localization precision is (FWHM) 61 nm for Z plane.

All measurements were performed on reconstructed super resolution images in Vutara SRX software. Diameter measurements were obtained by measuring the distance between the highest particle count located at the beginning of the structure and the highest particle count located at the end of the structure. Occasionally, we were unable to fully resolve the initial cortical rings. This is probably due to the depth of field limitations of the Vutara system (1 mm). In such cases, the diameter of the rings was determined based on the lateral dimension of the rings. For representation reasons, a fully resolved ring with a diameter of about 700 nm is presented in figure 1A. Scale bar is adapted accordingly.

## ACKNOWLEDGMENTS

We thank Michael Kozlov (TAU) for fruitful discussion on the mechanism of ESCRT-III mediated membrane constriction. We also thank Michael Elbaum (WIS) for access to the microscopy unit. Super-resolution microscopy was performed at the Irving and Cherna Moskowitz Center for Nano and Bio-Nano Imaging at the Weizmann Institute of Science. We thank Dikla Nachmias and all members of the Elia laboratory for technical help and support. This work is funded by the Israeli Science Foundation (ISF) Grant no. 455/13 and the Marie Curie Integration Grant (CIG).

## AUTHOR CONTRIBUTIONS

I.G designed and performed all experiments including imaging and data processing and analysis. T.D provided hands-on experience on the Vutara system, performed experiments and was involved in data processing. N.E. designed the experiments was involved in data analysis and interpretation and wrote the manuscript.

